# TMPs Interaction Domain: A Discovery of Structural Universality Among TMPs Interaction Sites Using Deep Learning Method

**DOI:** 10.1101/2021.05.19.444734

**Authors:** Yihang Bao, Weixi Wang, Minglong Dong, Fei He, Han Wang

## Abstract

Transmembrane proteins (TMPs) serve as important drug targets and accounts for nearly half of the drugs currently available in the market. Research into TMPs interactions and their structural basis will provide key information for drug research and new drug development. Based on previous works like a binding pocket or binding site, our main purpose in this study is to find whether the structural universality (Interaction Domain) exists in all kinds of TMPs interaction regions through a computational approach. After implementing the experiments using our 3D deep learning model and achieve the Matthews correlation coefficient (MCC) of 0.36, we found strong evidence for the existence of the structural basis among TMPs interaction regions. That means those regions, or we call them interaction domains, are structural specific distinguishing to the domains without any interaction. According to this, this work provides a new theoretical basis for TMPs interaction research and can greatly boost the development of the drug industry.

## 1 Introduction

Transmembrane Proteins (TMPs), as an essential type of Membrane Proteins(MPs), span the entire biological membrane with segments exposed to both inside and outside of the lipid bilayers(Stillwell 2016). Since TMPs play numerous roles in basic physiology and pathophysiology (Puder, Fischer, and Mierke 2019; He et al. 2019; Oguro and Imaoka 2019), they are major targets for nearly half of the known drugs on the current therapeutics market. In the field of drug development, for one thing, interaction information of TMPs helps researchers understand the drug interactions on a molecular biology level, leading the drug design more accurate and efficient(Lee et al. 2007). For another, structural information of interaction regions can greatly boost the new target screen and help researchers understand the drug-disease relations with better clarify(Koes and Camacho 2012). Unfortunately, since it is difficult to simulate the biofilm environment or just entice it a crystal, traditional experimental methods are ineffective to obtain the TMPs’ interaction information (Chen and Gingras 2007; McDowall, Scott, and Barton 2009). And that also results in the lack of known samples which blocks the bioinformatics approach utilized in soluble proteins widely. Meanwhile, existing studies just focused on their own partial research areas of protein interactions and the general basis among interaction regions has not been figured yet. All of this is calling for deeper research into the structural universality among TMPs interaction regions through bioinformatics methods.

Many previous efforts have been made in TMPs interaction research from various angles owing to its great importance. Some researchers explore TMPs interactions based on known structures. For example, Tusnady et al. extracted MPs’ interactive relationships from their topology structure(Dobson, Reményi, and Tusnády 2015) and Hurwitz et al. focused on MPs docking prediction relying on their 3D structure(Hurwitz, Schneidman-Duhovny, and Wolfson 2016). These researches need highly reliable known TMPs structure information but both bioinformatics methods and experimental methods are helpless to provide(Xiao and Shen 2015). Another group of researchers employed a biological network. Qi et al. took this kind of method to predict human membrane receptor interactions(Qi et al. 2009) but the false positive rate is high according to the lack of structural support. In consideration of these limitations, some researchers have started to conduct interaction studies from the perspective of structure feature mining. Kozlovskii et al. used the deep learning method to dig the structure of known protein-ligand complexes and achieved satisfactory results in certain druggable binding site identification(Kozlovskii and Popov 2020). It can be seen that all these works without exception encounter limitations so that they can only focused on a certain mode of interactions like docking and binding or certain interaction materials like metal ions, small molecules and ligands. They all tried to predict the certain interaction information directly, but ignore the structural universality among all kinds of interaction sites and the potential help of structural universality to the prediction of interaction information.

Based on the knowledge that protein structure determines function(Orengo, Todd, and Thornton 1999), we deem the local interaction domain as the direct structural basis of TMPs’ interaction function. Consists of a group of adjacent residues near the protein surface, TMPs interaction domain has structural stability and specificity that can determine certain interaction activities. The interaction domain here is a collection of various interaction modes including docking and binding. After obtaining a series of interaction domain information, we can make use of relevant sequence-based features to efficiently predict specific interaction information. Figure 1 shows our idea. This idea breaks through the existing barriers and develops a new feasible direction for TMPs interaction research. Firstly, the dependence on known TMPs structure and interaction relationships is reduced. Secondly, the interaction domain as a whole will have stronger sequence feature significance, and the feature learning for a group of TMPs surface residues will obviously be better than the feature learning for a single residue due to the increase of information. Furthermore, after the problem of TMPs interaction prediction is transformed into the problem of TMPs interaction domain discovery, different types of interaction materials can be modeled in a unified way. Whereas there have been no previous studies on the TMPs interaction domain, the existence of TMPs interaction domain needs to be verified urgently, or to be more specific, we need to verify whether the structural universality among TMPs interaction regions exists. This problem of learning features directly from spatial structures has puzzled researchers for years due to the limitations of the algorithm and hardware. Fortunately, since Qi et al. first proposed Pointnet(Qi, Su, et al. 2017) and Pointnet++(Qi, Yi, et al. 2017), abundant deep learning method based on CNN(Komarichev, Zhong, and Hua 2019) or GNN(Xu et al. 2020) were proposed to directly learn deep features from the raw three-dimensional point cloud data. These methods have also been adopted in bioinformatics like Mallet et al. tried to discover the relationship between ligand and protein structure using Voxel CNN(Mallet et al. 2019). All of this demonstrates that 3D learning methods are able to learn deep features from protein spatial structure efficiently.

**Figure 1.**
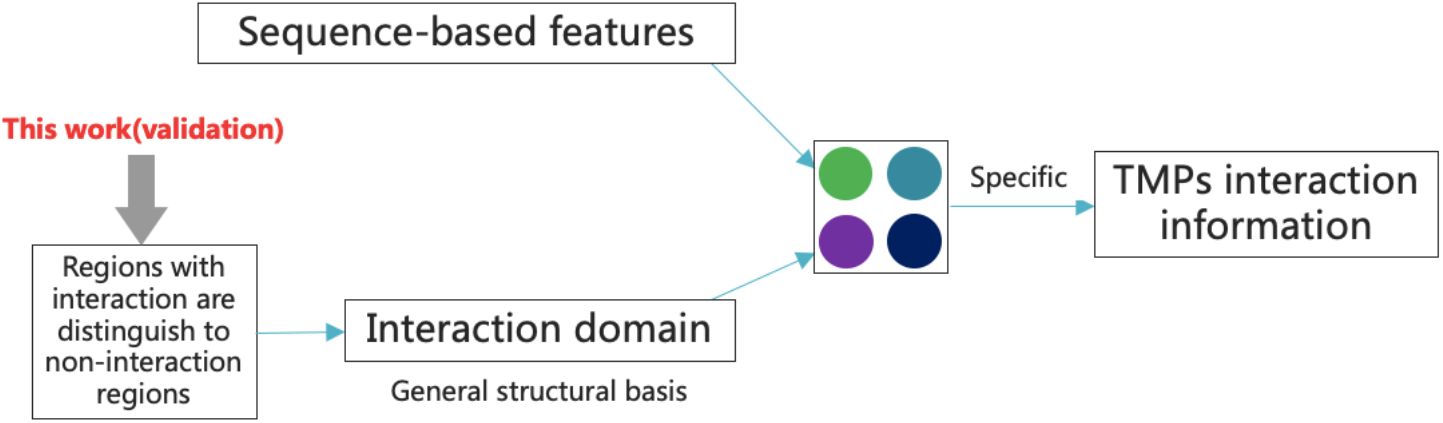
Our new idea to explore the TMPs interaction information

In this study, in order to study the TMPs related interaction activities from a more general perspective, we proposed the conjecture of TMPs Interaction Domain and tried to certify the existence of this structural universality among all kinds of TMPs interaction regions using the 3D deep learning method. The whole proof process is mainly composed of two parts, namely statistical analysis and prediction of TMPs interaction domain. In the statistical analysis part, since there is no previous works indicate the exact universality among all kinds of TMPs interaction regions as well as the size of the interaction domain, we need to determine it statistically from known 3D structures of the existing Multi-Residue-Interaction-Sites (MRIS). After collecting data from Protein Data Bank (PDB) (Berman et al. 2000), we determine the preliminary feature and the sampling window of the interaction site to make it suitable for most of the interaction regions. And in the prediction part, the fundamental purpose is to verify the existence of general features. More specifically, once our deep learning model is capable of satisfactorily classify whether or not a discretionary group of residues constitutes a TMPs interaction domain after training, we can say that this kind of universality (Interaction Domain) exists. For this purpose, we adopted an end-to-end learning network modified from Pointnet(Qi, Su, et al. 2017). Equipped with a robust network, our model efficiently reached the Matthew Correlation Coefficient (MCC) of 0.36 on an independent dataset with the actual ratio of positive samples and negative samples. This wonderful MCC not only lends support to our conjecture, but also indicated that this whole new angle for protein interaction research and drug development is feasible and has great potentiality. Both pre-trained model and supported information can be found at https://github.com/NENUBioCompute/TMPInteractionDomain

## 2 Materials

As mentioned above, 3D structure information of the TMPs interaction sites is required to construct our dataset. According to this, 4313 TMPs were downloaded from Protein Data Bank (PDB)(Berman et al. 2000), the biggest protein 3D information library, with the PDBID found in the Protein Data Bank of transmembrane proteins (PDBTM)(Kozma, Simon, and Tusnady 2012). We did not directly download them from PDBTM because files in PDBTM may be obsolete or faulty.

By utilizing the Biopython tools(Cock et al. 2009) and PDBparser module developed by ourselves, we reformatted the PDB files and extracted site information from them. And for feature analysis, we also did the similar in other regions. The data format is shown in **Table 1**.

**Table 1.** Raw data format

| Field | Meaning |
| --- | --- |
| SiteID | Name of the interaction site |
| numRes | Number of residues |
| EVIDENCE_CODE | From remark 800 of PDB |
| SITE_DESCRIPTION | From remark 800 of PDB, the conjugate |
| center_res_ca_coord | Coordinate of the closest alpha carbon atom to the center |
| center_res_ca_depth | Depth of the closest alpha carbon atom to the center |
| res_info | Information of every single residues |

## 3 Feature Analysis and Spatial Scale Determination through Statistical Analysis

### 3.1 Preliminary Exploration of Universal Features

Since there is no previous work that proved the existence of structure universality in all kinds of TMPs interaction religions, we tried to explore it based on the ground truth data from PDB. We randomly extracted residues from PDB files of 1000 TMPs. For each residue, we counted the average distance of the nearest N residue and classified the average distance according to the number of interaction site residues contained in these N+1 residues. The result can be seen in **Figure1**. It can be found that with the increase of the number of residues belonging to interaction sites in the surrounding space, the average distance between randomly selected residues and their nearest n residues tends to increase, which means space tends to be sparser. This ground truth indicates that the space near the interaction regions does have some spatial features that are different from other non-interaction regions. We will refer to this specific region as the interaction domain in the following paragraphs.

### 3.2 Relative Residual Depth

Although most of the TMPs interaction domains are on the solution accessible surface, there still exist some TMPs interaction domain hidden inside. Such domains may expose on the surface after proteins have been unfolded or bent(Tan et al. 2013). According to this, we modified the concept of residual depth, the distance between residues and accessible surface(Chakravarty and Varadarajan 1999), and proposed relative residual depth. After choosing the central residue from all residues in the interaction site as K, the relative depth of the remaining residue is the absolute value of the difference from K. In consideration of the inside sites, this computing method is more convincing to measure the exact scale of the TMPs interaction domain. **Table 2** shows the relationship between residue coverage rate and relative residual depth in interaction sites with a certain number of residues. Residue coverage here is defined as the number of residues less than the relative residue depth divided by the total number of residues.

**Table 2.** Relative residue depth statistics in interaction sites with certain number of residues

| Number of residues<br>each site | $\leq 3\text{\AA}$ | $\leq 4\text{\AA}$ | $\leq 5\text{\AA}$ | $\leq 6\text{\AA}$ |
| --- | --- | --- | --- | --- |
| 2 | 0.948 | 0.976 | 0.988 | 0.993 |
| 4 | 0.881 | 0.946 | 0.973 | 0.987 |
| 6 | 0.834 | 0.922 | 0.963 | 0.986 |
| 8 | 0.786 | 0.896 | 0.945 | 0.970 |
| 10 | 0.728 | 0.849 | 0.914 | 0.951 |
| 12 | 0.704 | 0.824 | 0.892 | 0.934 |
| 14 | 0.684 | 0.798 | 0.872 | 0.923 |

### 3.3 Sampling Window of Protein Interaction Domain

Sampling window is a set of sampling principles and has been utilized in plenty of machine learning tasks(Salle, Idiart, and Villavicencio 2016). In this part, we try to take a given number of residues in a given range from the center and the mission of our sampling window is to determine this number and range based on the coverage rate and biological significances. The number, or we call it window width, should be approximately between 15 and 35 after analyzing the data features. The range, or we call it window depth, is the relative residue depth mentioned above and its value should be approximately between 6 and 8 based on Table 2. These two values are combined to form the sampling window and the residue coverage at different values can be seen in **Table 3**.

**Table 3.**
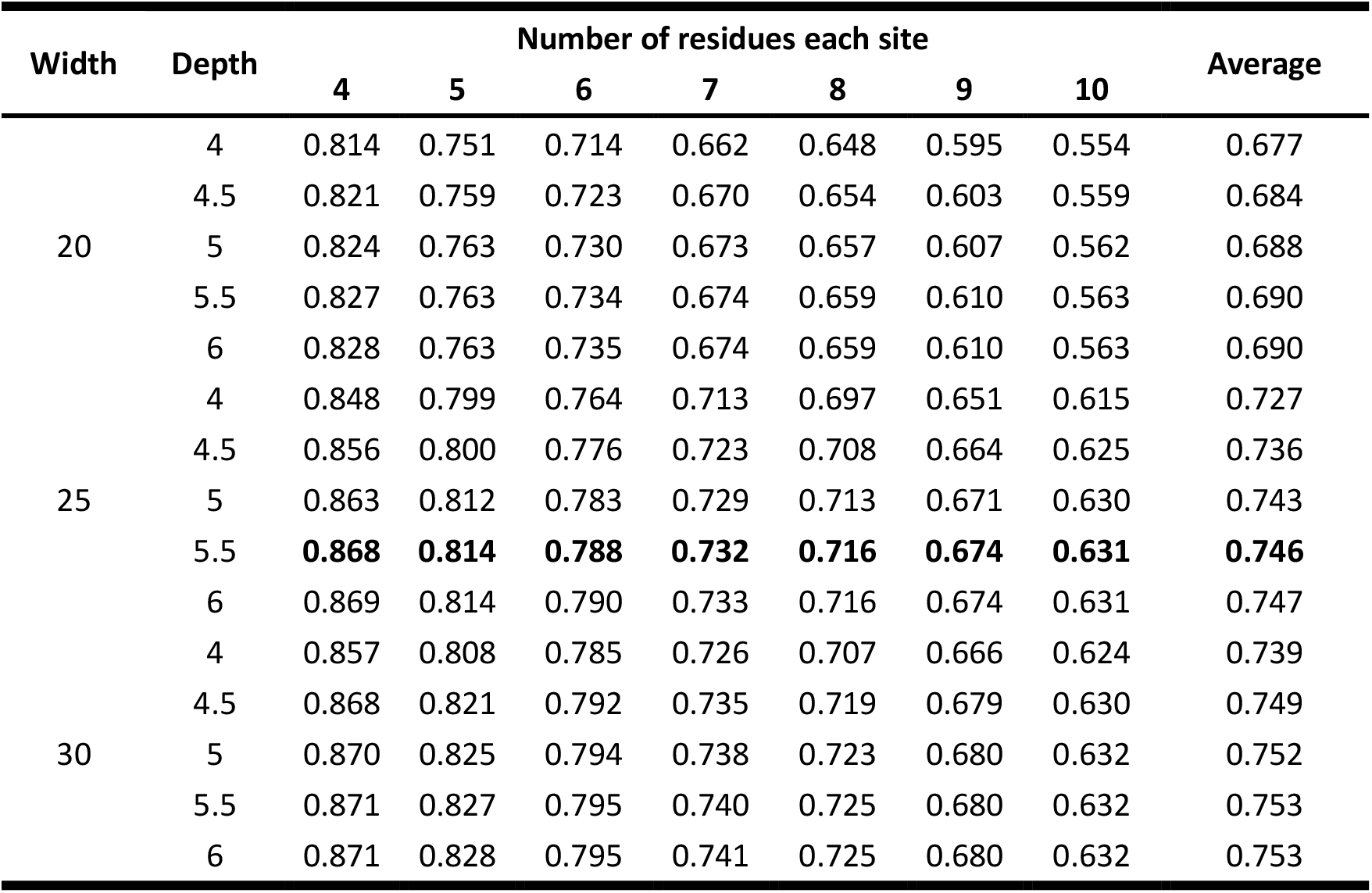
Residue coverage with different sampling window

Obviously, the residue coverage increases monotonically as window width or depth increases, but that does not mean the higher the residue coverage the better the performance. Once the sampling window we take is much larger than the actual interaction domain of lots of sites, this kind of redundancy will significantly reduce its structure specificity and waste computing resources(Redlich 1993). It is as if the smaller an apple is in the picture, the harder it is to identify it from the picture. According to this, though the residue coverage rate is highest at width 30 and depth 6, the increase in coverage is very limited compared to some smaller parameters. And meanwhile, they pay the price of significantly higher redundancy. Therefore, we adopt width 25 and height 5.5 as the parameters of our sampling window. This sampling window covers the vast majority of interaction domains, with the usability and representativeness of pattern learning.

### 3.4 Data Sampling and Partitioning

In machine learning tasks, we need positive and negative samples(Michie, Spiegelhalter, and Taylor 1994). For positive samples, the sampling window parameters mentioned above are adopted. To be more specific, we selected 25 residues near each site as positive samples of the interaction domain and did not sample when the number of residues exceeded 25 or the relative residue depth exceeded 5.5. For negative samples, we traversed all residues that are not in the positive samples and sampled with α carbon atoms of each residue as the center. If there existed residues in the positive sample after sampling, they will be discarded. After removing samples with high similarity, we finally have 37961 positive samples and 3598175 negative samples. After the positive and negative samples were combined and randomly shuffled, the whole dataset was randomly divided into the training set, validation set and test set according to the ratio of 7:2:1. It is worth noting that the positive and negative sample ratios in the test set and validation set are consistent with the real situation, leading our evaluation results more realistic and convincing.

## 4 Model Design and Implementation

### 4.1 Features and Input Encodings

Features are the key issue for the machine learning tasks(Zhang and Liu 2019). In this study, One-hot coding(Cassel and Lima 2006) is adopted to encode the types of residues and three-dimensional coordinates were used to characterize the spatial features of residues(See **Figure 1**). Since what we need to prove is structural universality, the three-dimensional coordinates are necessary and significant for spatial features description(Senior et al. 2020). Meanwhile, once combine the coordinate information and the residue type information, more physical and chemical features can be obtained(Morris et al. 1992).

### 4.2 Network Architecture

Inspired by the ground truth explored in section 3.1, here we tried to construct our model based on classic point cloud deep learning models. As a deep learning method, our model can predict whether a discretionary group of residues constitutes a TMPs interaction domain. As we can see in **Figure 2**, the model was modified from Pointnet(Qi, Yi, et al. 2017) and added a subnet structure for inputting amino acid names and voting mechanism. The whole model can be divided into 4 parts, which are coordinates processing, residue name processing, classification, and voting. The coordinates processing part was used to extract deep features of 3D coordinates, which is similar to the operation of PointNet. And in consideration of the consistency of processing, we also adopted STN, weight-shared MLP and max-pooling(Nagi et al. 2011) to extract deep features from One-hot code in the residue name processing part. Once we got these two deep features, we connected them into a global feature and fed them into two fully connected hidden layers with a sigmoid-activated output layer, which outputted the binary result.

**Figure 2.**
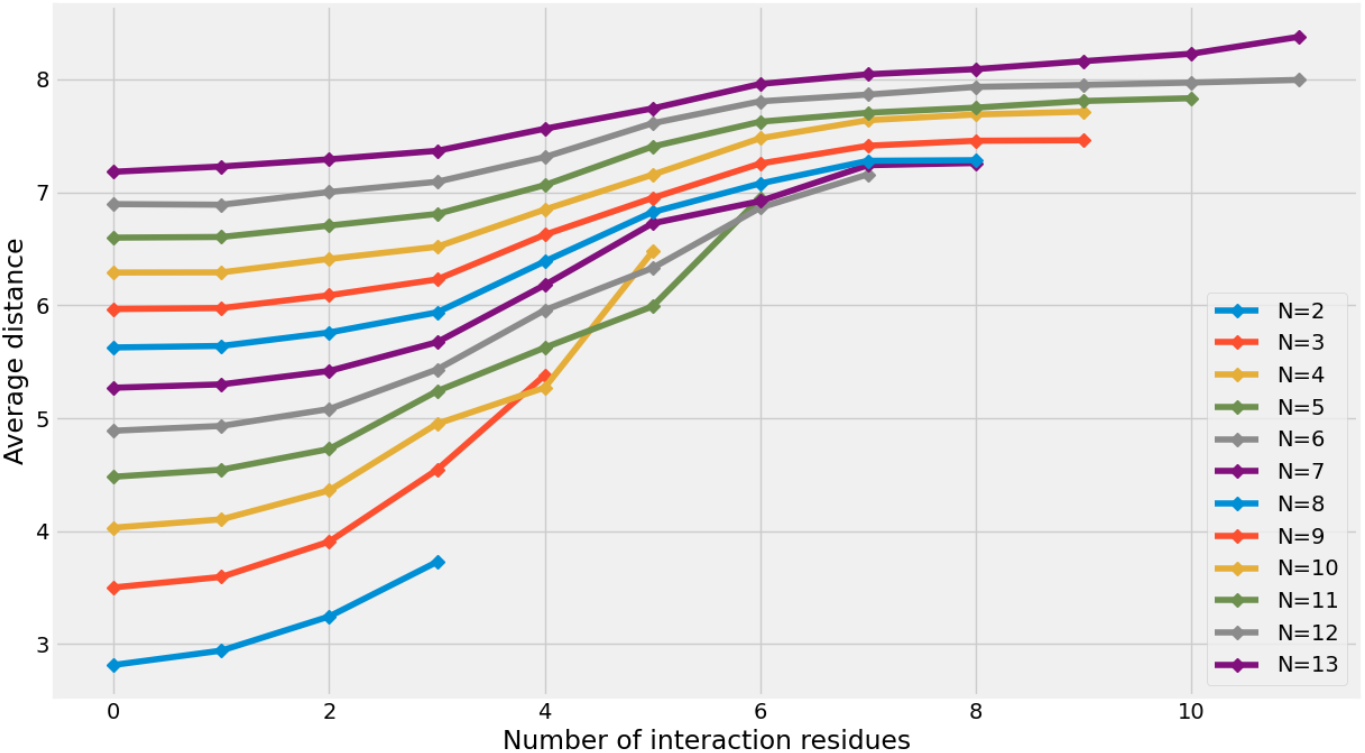
Average distance between randomly selected residues and their nearest n residues

As a binary classification problem, we utilized BinaryCrossentropy as the lost function (Creswell, Arulkumaran, and Bharath 2017) and Adam as the optimizer(Zhang 2018). Besides, in order to solve the problem of the unbalanced dataset, we added an extra voting part to our model. After each model had been trained and optimized separately on its own training set, we vote on the result of these models to get the final result. More details about the voting mechanism are discussed in section5.

### 4.3 Performance Evaluation

An intuitive evaluation index for deep learning results is accuracy, which is the proportion of the total number of correct predictions. Nevertheless, accuracy is meaningless here because our data set is extremely unbalanced(Yu, Chou, and Chang 2010). To quantitatively evaluate the performance, we adopted six measures including accuracy, recall, precision, specificity, MCC and F-measure(Tan et al. 2019; Yang et al. 2019; Zhu et al. 2019). Precision, recall, and specificity describe the data from different perspectives while MCC and F-measure are composite indexes (See **Table4**.).

**Table 4.** Description of evaluation indexes

| Index | Formulas | Description |
| --- | --- | --- |
| Recall | $\frac{TP}{TP + FN}$ | The proportion of predicted interaction domain to the total number of the interaction domain |
| Precision | $\frac{TP}{TP + FP}$ | How likely is it that the predicted positive sample is actually an interaction domain |
| Specificity | $\frac{TN}{FP + TN}$ | How likely is it that the predicted negative sample is not an interaction domain |
| MCC | $\frac{TP \times TN - FP \times FN}{\sqrt{(TP + FP)(TP + FN)(TN + FP)(TN + FN)}}$ | Index to measure the overall performance of the binary predictor |
| F-measure | $2 \times \frac{Recall \times Precision}{Recall + Precision}$ | Comprehensive consideration Recall and Precision |

where TN, TP, FN, and FP, respectively denoted true negative, true positive, false negative, and false positive

### 4.4 Voting Mechanism Analysis

The parameter of the voting mechanism is a kind of threshold, or in other words, once the number of positive voters exceeds this threshold, the result is positive. **Table 5** shows the performance of our model on the validation dataset under different voting mechanism parameters. As we can see that MCC on the validation dataset reached its maximum when the voting mechanism threshold is around 23. The voting mechanism here, in essence, is another implementation idea of over-sampling and under-sampling(Chawla et al. 2002; Drummond and Holte 2003) that is widely used in imbalanced dataset processing. And if we combine all 30 training sets, it is a typical over-sampling method. Compared with over-sampling and under-sampling, the voting mechanism is more flexible and effective at dealing with the extremely unbalanced dataset in this study.

**Table 5.** Performance on validation set under different voting mechanism threshold

| Voting mechanism threshold | MCC |
| --- | --- |
| 19 | 0.343 |
| 20 | 0.347 |
| 21 | 0.347 |
| 22 | 0.348 |
| 23 | **0.358** |
| 24 | 0.357 |
| 25 | 0.354 |
| 26 | 0.348 |

### 4.5 Implementation details

Our model was implemented, trained and tested adopting the popular deep learning library PyTorch(Paszke et al. 2019) on four Nvidia Tesla V100 GPUs. We set the batch size to 32 and the initial learning rate to 0.0001. The training dataset was divided into 30 equal parts, each of them had the same positive samples but different negative samples. Utilizing five-fold validation(Walsh, Pollastri, and Tosatto 2016), all training sets are trained independently, and their epochs are selected according to the MCC performance on the validation set. That is to say, 30 models trained on the training set may have different epochs. Actually, we have experimented with a lot of other 3D point cloud models like Grid-GCN(Xu et al. 2020) or Geo-CNN(Lan et al. 2019), but their models had no significant result improvement and brought huge computational overhead. This is because those models all focus on the processing of large-scale point clouds while not suitable for feature extraction of the small-scale point clouds in this study.

## 5 Results and Discussion

### 5.1 Overview of TMPs Interaction Domains in Test Set

The test set is randomly selected and has a real positive-negative ratio. Since it contains most of the interactions materials as well as interaction forms, this data set is general and can fully evaluate our model and conclusions. In the analysis of the test set, we found that the spatial distribution of the interaction domains may be related to the interaction materials. For example, metal ions such as copper prefer to interact in the internal region of the protein, while larger molecules prefer to interact on a relatively flat surface. In addition, approximately 40 percent of the interaction domain related spatial structures in the test set were identified by DrugBank(Wishart et al. 2018) as being closely related to drug targets. It indicates that the interaction domains have strong functional characteristics and have great potential in boosting the process of drug discovery.

### 5.2 Prediction Performance

**Table 6** shows the accuracy, recall, precision, specificity, MCC and F-measure result of our model on the test dataset using voting mechanism threshold 23. These results are presented separately based on family, interaction materials and drug target. As we can see that our model reaches an overall precision of 0.81. That means once our model considers a local space to be an interaction domain, it is very likely to be an interaction domain. This result effectively demonstrates that our model can classify interaction domain from dataset efficiently, or further, this result indicates that our model can classify these data with the help of the structural universality among TMPs interaction sites. Besides, our model can achieve similar evaluation indexes under the conditions of different types of data sets, which proves that what we have found is indeed a general spatial structure universality, rather than a universality for a specific case. Though the recall is less than satisfactory, that does not mean our hypothesis is doubtful. It just indicates that the dataset is not comprehensive enough or the way we processed the data, including feature selection or data sampling, still needs to be promoted and optimized.

**Table 6(A).**
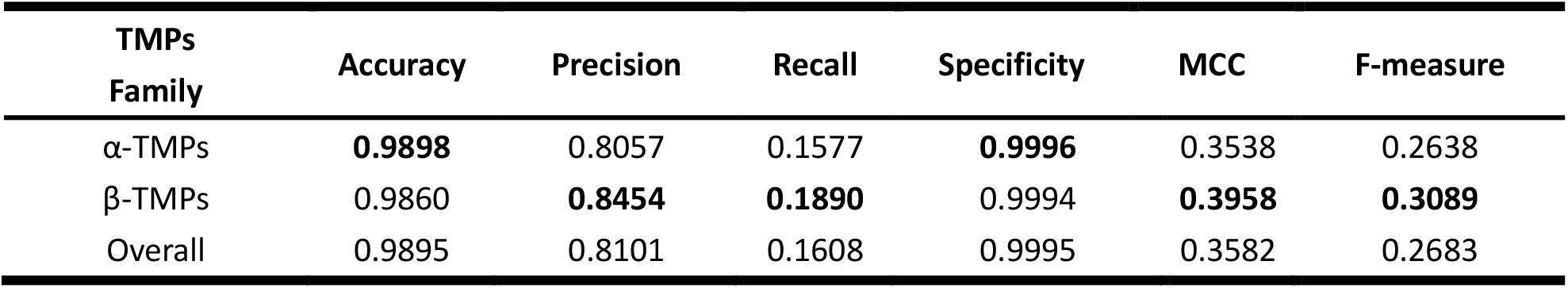
Model evaluation on the test set according to TMPs families

| TMPs Family | Accuracy | Precision | Recall | Specificity | MCC | F-measure |
| --- | --- | --- | --- | --- | --- | --- |
| α-TMPs | **0.9898** | 0.8057 | 0.1577 | **0.9996** | 0.3538 | 0.2638 |
| β-TMPs | 0.9860 | **0.8454** | **0.1890** | 0.9994 | **0.3958** | **0.3089** |
| Overall | 0.9895 | 0.8101 | 0.1608 | 0.9995 | 0.3582 | 0.2683 |

**Table 6(B).** Model evaluation on test set according to interaction materials

| Interaction Materials | Accuracy | Precision | Recall | Specificity | MCC | F-measure |
| --- | --- | --- | --- | --- | --- | --- |
| Saccharides | 0.9895 | 0.8102 | 0.1608 | 0.9995 | 0.3582 | 0.2684 |
| Peptide | **0.9896** | 0.8121 | 0.1614 | 0.9995 | **0.3593** | **0.2693** |
| Metal ion | 0.9895 | 0.8097 | 0.1604 | 0.9995 | 0.3576 | 0.2677 |
| RNA | **0.9896** | 0.8114 | **0.1610** | 0.9995 | 0.3585 | 0.2685 |
| Other small organic molecules | **0.9896** | **0.8123** | 0.1609 | **0.9996** | 0.3588 | 0.2686 |
| Others | 0.9895 | 0.8114 | 0.1607 | 0.9995 | 0.3584 | 0.2683 |
| Overall | 0.9895 | 0.8101 | 0.1608 | 0.9995 | 0.3582 | 0.2683 |

**Table 6(C).** Model evaluation on test set according to whether it is a drug target

| Drug Target | Accuracy | Precision | Recall | Specificity | MCC | F-measure |
| --- | --- | --- | --- | --- | --- | --- |
| Yes | 0.9895 | 0.8097 | 0.1601 | 0.9995 | 0.3573 | 0.2673 |
| No | 0.9895 | **0.8117** | **0.1617** | 0.9995 | **0.3595** | **0.2696** |
| Overall | 0.9895 | 0.8101 | 0.1608 | 0.9995 | 0.3582 | 0.2683 |

### 5.3 Case Study

In order to further demonstrate the existence of TMPs interaction domain, we randomly took Undecyl-Maltopyranoside(UMQ), Protoporphyrin IX Containing FE(HEM) and HEME C(HEC) as example interaction materials of our case study. UMQ is a kind of small molecule and can affect hydrogen ion transmembrane transporter activity through target V-type sodium ATPase subunit K(Overington, Al-Lazikani, and Hopkins 2006). HEM and HEC are structurally similar while HEC plays vital roles in lots of protein-based activities such as metal ion binding based on target cytochrome c4 and iron ion binding based on target high-molecular-weight cytochrome(Imming, Sinning, and Meyer 2006). After screening related and correctly predicted interaction domains from our test dataset, they are visualized in **Figure 4** using PyMOL(DeLano 2002). Blue parts are predicted interaction domains while red parts are interaction materials. This picture assists to draw the following conclusions. Firstly, our model works. Interaction domains in the first row of Figure 4 are extracted from highly homologous proteins and our model is able to locate them accurately from the different spatial coordinate systems. Secondly, diverse sequences can form the similar interaction domain for a particular interaction material. By comparing the subfigure in the second or third row of Figure 4, it can be seen that different sequences can form similar spatial structures in space to interact with the same materials. Finally, similar spatial structures can interact with different materials. The second and third row of Figure 4 have the similar spatial structures of the interaction domain, but they interact with different kinds of materials.

**Figure 3.**
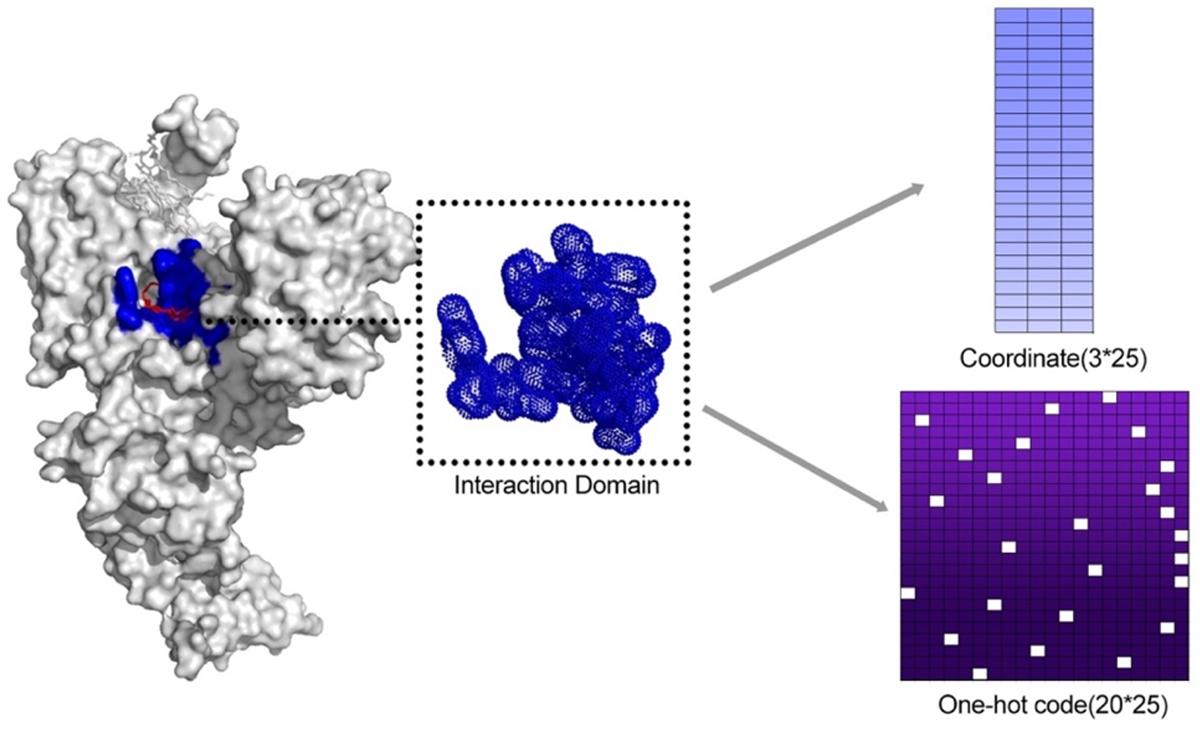
Coordinates and One-hot code of residue name are extracted to represent the interaction domain

**Figure 4.**
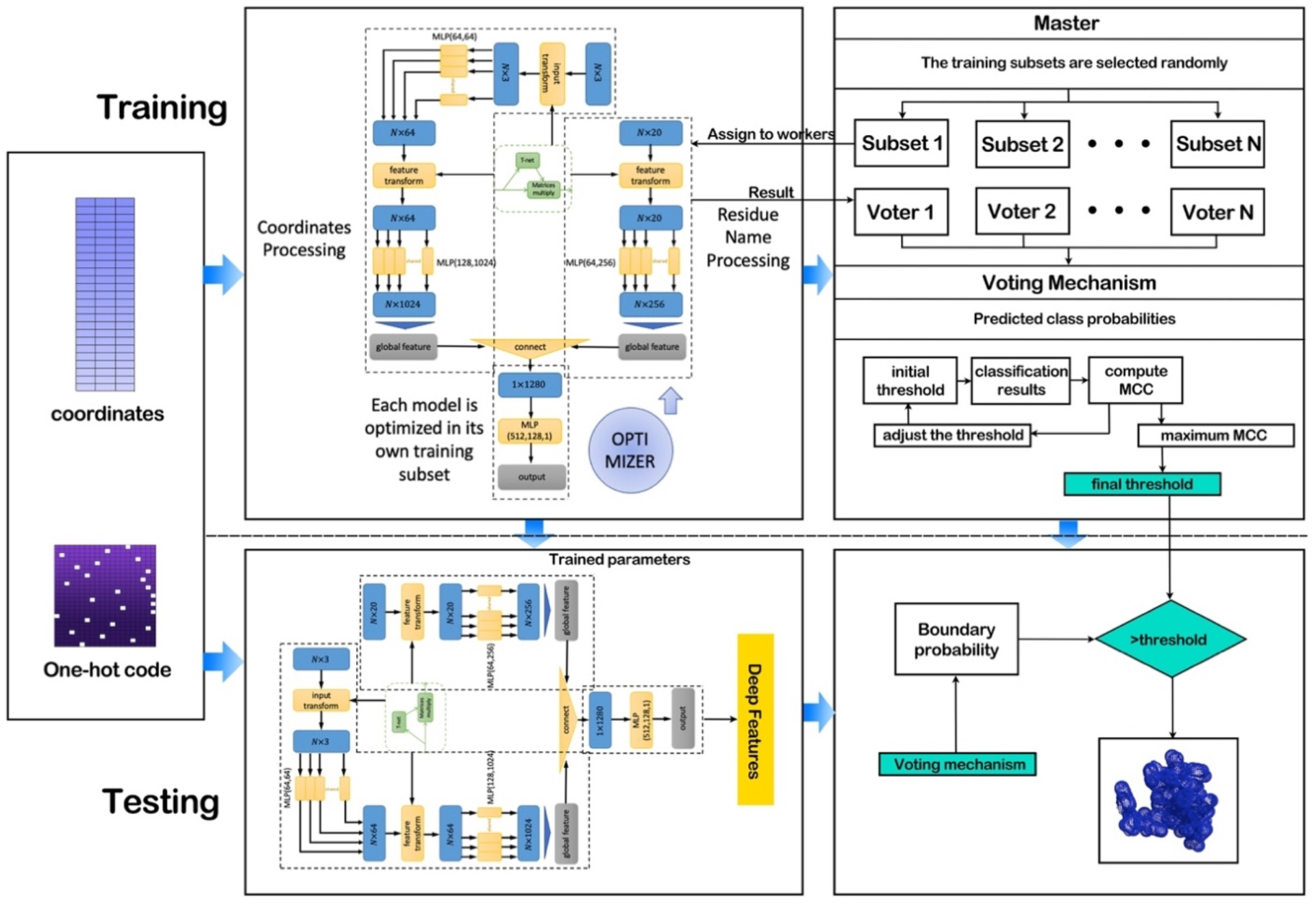
Network architecture

In addition, after revisiting the non-interaction samples that our model predicted as positive, we found that some of them were actually positive samples but are not listed in the PDB. Figure 6 shows the interaction between Nevirapine and protein 1JLB where the information is provided by BindingDB(Liu et al. 2007). Nevirapine is a potent, non-nucleoside reverse transcriptase inhibitor (NNRTI) used in combination with nucleoside analogues for the treatment of Human Immunodeficiency Virus Type 1 (HIV-1) infection and AIDS(Wishart et al. 2018). This indicates that the dataset we used in training is defective as the positive samples are hard to find through experimental methods. That is, the unsatisfactory model performance is mostly not due to the poor model learning ability or lack of general structural universality. We believe that this kind of general structural universality will be more significant with more precise datasets.

**Figure 5.**
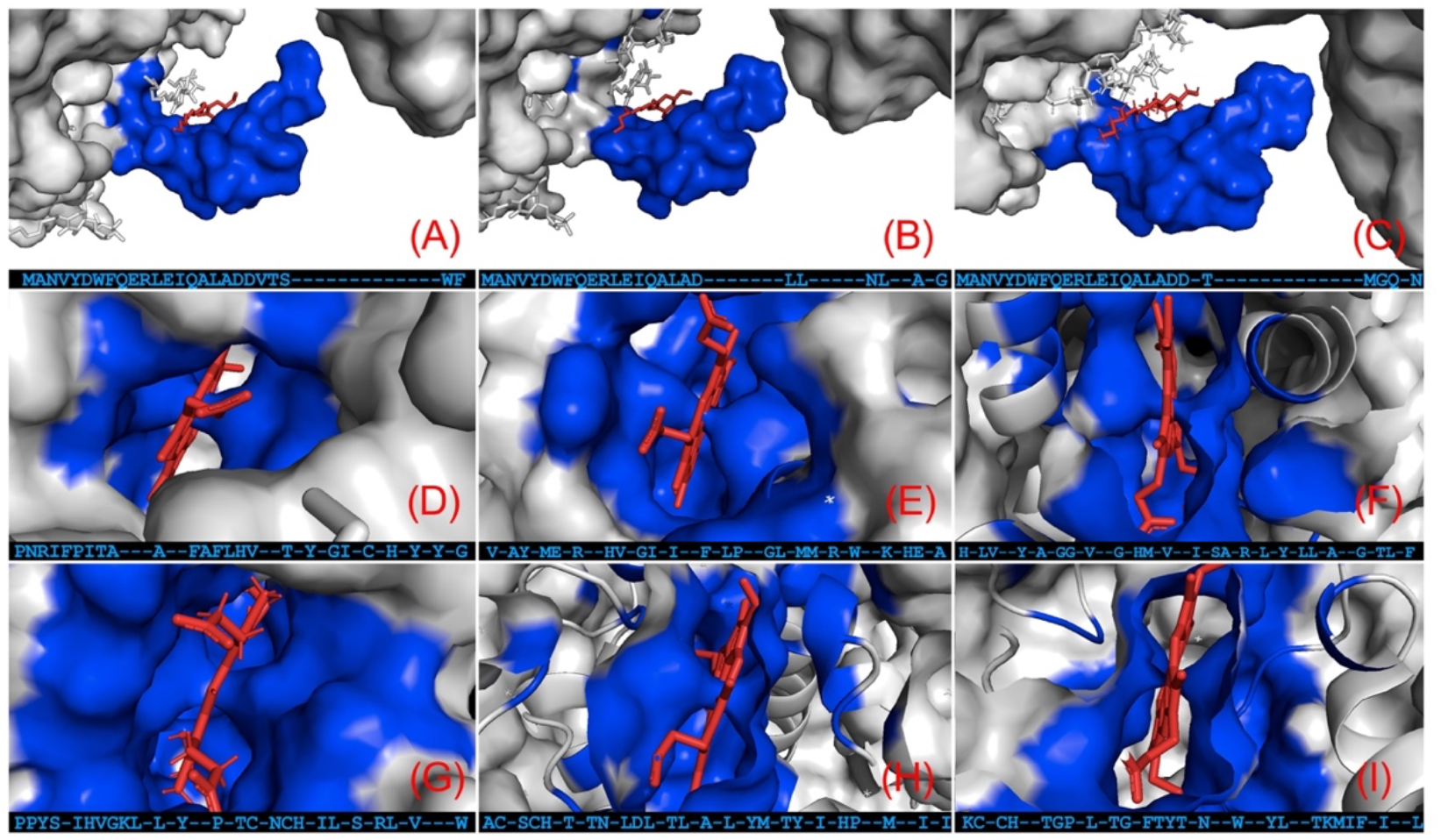
(A)(B)(C) UMQ interaction domains in proteins with PDB ID of 2E74, 2E76, 4H13; These three pictures illustrate that our model is effective in homologous proteins (D)(E)(F) HEM interaction domains in proteins with PDB ID of 4H0L, 5OC0, 5XMJ; These three pictures illustrate that diverse sequences can form the similar interaction domain (G)(H)(I) HEC interaction domains in protein with PDB ID of 6F0K, 5B5E, 4HL3; These three pictures illustrate that similar spatial structures can interact with different materials

**Figure 6.**
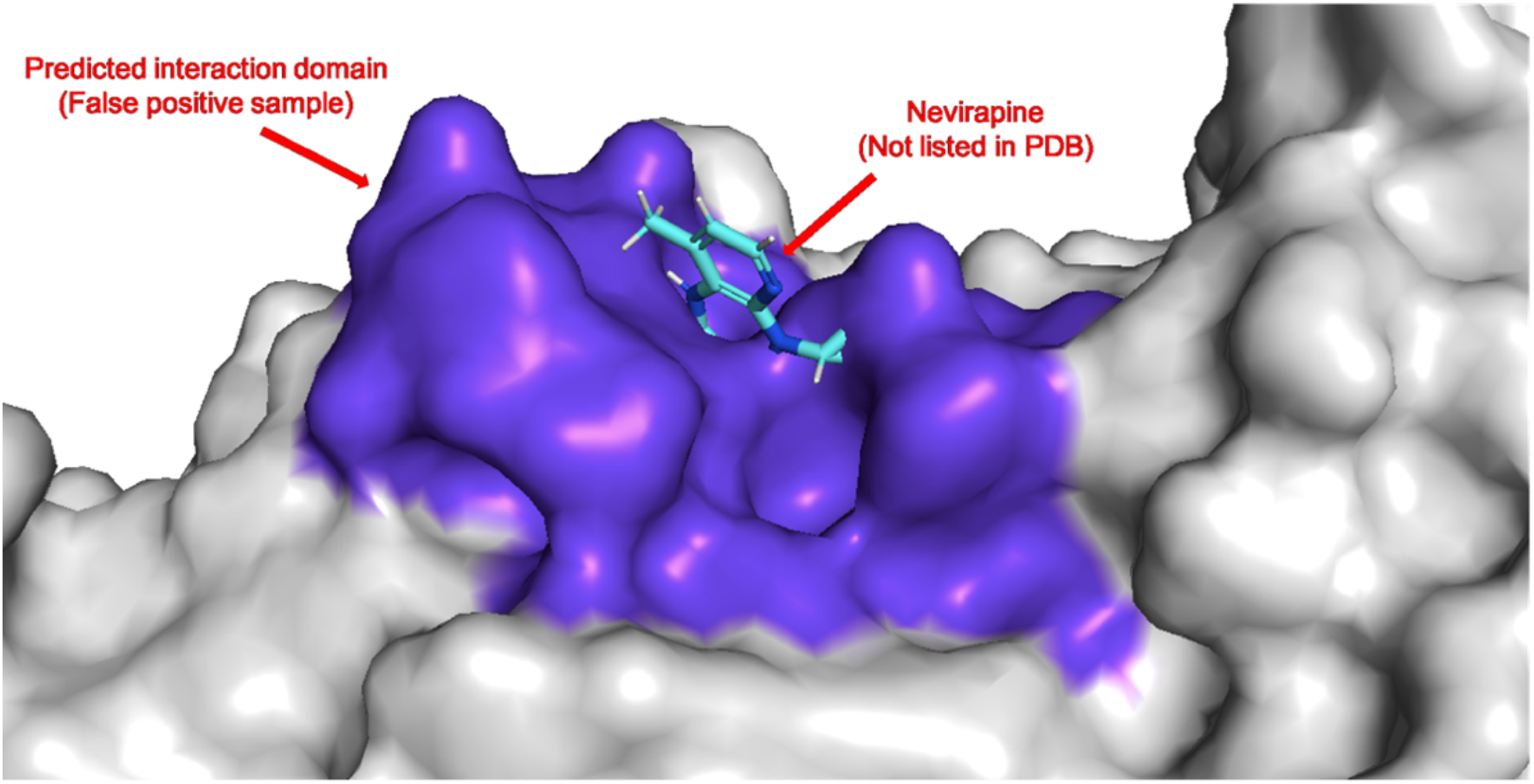
Part of the false-positive samples (e.g. interaction between Nevirapine and protein 1JLB) are discovered as interaction domain according to external databases

## 6 Conclusions

In this study, we proposed the conjecture of TMPs Interaction Domain and tried to certify this structural universality among TMPs interaction sites using the 3D deep learning method. Modified from PointNet, our model combines the 3D coordinates and residue to extract deep features from the protein’s local structure and adopt a voting mechanism to solve the problem of the unbalanced dataset. Though some of the data processing methods need to be improved, the MCC performance of this model on the test set still reached 0.36. Subsequently, we analyzed our prediction result using PyMOL as strong evidence to prove the existence of structural universality.

According to this, we proved the existence of interaction domain from both model output and case study result. Based on this, we will develop a more accurate prediction model of interaction domains using more efficient data processing methods and learning models. As long as we get a spatial structure that is more likely to interact, we can further determine the specific interaction mode and interaction materials of this spatial structure with the help of sequence-based features. This two-step prediction method is more effective than the traditional method. Firstly, our methods are less dependent on the sample size of known membrane protein structures and interaction relationships. Secondly, stronger sequence feature significance will be represented. Besides, we can predict all kinds of interactions in a unified way.

In conclusion, this finding of TMPs interaction domain not only provides the feasibility for effectively predicting membrane protein interactions, but also opens a new research angle for drug development and many other works. We hope our work truly helps.

## Conflicts of Interest

All authors affirmed that there are no conflicts of interest

## Author Contributions

YB and WW conceived the idea of this research, processed data, and implemented the model. YB tuned the model and wrote the manuscript. HW and FH supervised the research, gave guidance during the research, and reviewed the manuscript. All authors were involved in the work and approved the submitted version.

## Funding

This work was supported by the Jilin Scientific and Technological Development Program (No. 20180414006GH), and Fundamental Research Funds for the Central Universities (Nos. 2412019FZ052 and 2412019FZ048).

## Notes

https://github.com/NENUBioCompute/TMPInteractionDomain

### Competing Interest Statement

The authors have declared no competing interest.

